# NEMO: Improved and accurate models for identification of 6mA using Nanopore sequencing

**DOI:** 10.1101/2024.03.12.584205

**Authors:** Onkar Kulkarni, Lamuk Zaveri, Reuben Jacob Mathew, Nitesh Kumar Singh, Sreenivas Ara, Shambhavi Garde, Manjula Reddy, Karthik Bharadwaj Tallapaka, Divya Tej Sowpati

## Abstract

DNA methylation plays a key role in epigenetic regulation across lifeforms. Nanopore sequencing enables direct detection of base modifications. While multiple tools are currently available for studying 5-methylcytosine (5mC), there is a paucity of models that can detect 6-methyladenine (6mA) from raw nanopore data. Leveraging the motif-driven nature of bacterial methylation systems, we generated 6mA identification models that vastly surpass the accuracy of the current best model. Our work enables the study of 6mA at a single-base resolution in new as well as existing nanopore datasets.

## Background

DNA methylation is a fundamental mechanism that regulates various biological processes, including embryonic development, genomic imprinting, X-chromosome inactivation, and suppression of transposable elements^1–3^. The methylation status of DNA is highly dynamic and can change in response to various environmental cues, such as stress, diet, and aging^4,5^. Aberrant DNA methylation patterns have been associated with various diseases, including cancer, neurodegenerative disorders, and cardiovascular diseases^6,7^. Moreover, recent studies have shown that DNA methylation patterns can be used as diagnostic and prognostic markers for various diseases and as potential therapeutic targets^8^. The most commonly studied methylated bases of DNA are 5-methylcytosine (5mC), 5-hydroxy methylcytosine (5hmC), and 6-methyladenine.

6-methyladenine (6mA) is an epigenetic modification of DNA that has long been known to exist in prokaryotes, where it is involved in mechanisms such as DNA repair and protection against foreign DNA^9,10^. Initially thought to be absent in eukaryotes, recent studies have shown that 6mA is present in several eukaryotic genomes, including fungi, plants, and animals^11–13^. In eukaryotes, 6mA is involved in several biological processes, including gene regulation, DNA replication, and chromatin structure, suggesting that this modification plays a crucial role in epigenetic regulation^11^.

Unlike 5mC, identification of 6mA is not amenable to conversion based methods such as bisulfite sequencing. Traditionally, 6mA is studied using capture based methods such as methylated DNA immunoprecipitation (MeDIP), where methylated DNA is enriched using antibodies specific to 6mA followed by high-throughput sequencing^14^. Such methods provide genome wide data on 6mA patterns, but lack nucleotide-level resolution. Recently, emerging single molecule sequencing technologies such as PacBio and Oxford Nanopore (ONT) sequencing have enabled direct identification of DNA base modifications.

ONT sequencing uses a voltage sensor to monitor electrical fluctuations as a DNA strand is translocated through a biological nanopore^15^. These characteristic fluctuations are converted to the corresponding DNA sequence by a basecaller. The ability to detect DNA modifications is due to the differences in the signal when a modified base translocates through the nanopore as compared to a canonical base^16^. Leveraging this, several tools have been developed that identify base modifications from raw nanopore data. A large subset of these are focused on 5mC, particularly in the CpG context^17^. A few tools support 6mA calling: Tombo, DeepSignal, DeepMP, and mCaller. Tombo, an old tool provided by Oxford Nanopore, identifies possible modified bases by testing statistically significant differences in the current levels among reads aligning to the target position^18^. DeepSignal and DeepMP are two recent Deep Neural Network based tools which provide models for 6mA identification only in the GATC context ^19,20^. mCaller provides a “dinucleotide” model, which theoretically covers all possible dimers with 6mA in the 1st position^21^. The performance of these tools in diverse sequence contexts remains unknown due to lack of appropriate benchmarking datasets.

In this study, we set out to develop accurate models that can discriminate 6mA from a canonical adenine from raw nanopore signal data generated on the R9.4.1 chemistry. To achieve this, we utilized datasets from multiple bacterial species for which sequence context of 6mA is known. We call our models NEMO (Nanopore Epigenetic Modification Output). We show that NEMO models significantly outperform existing models in identification of 6mA, both in GATC as well as other sequence contexts.

## Results

### Dataset generation and validation

Most of the existing 6mA callers have been trained and validated on limited datasets, such as a pUC plasmid grown in presence of *Escherichia coli* dam (DNA adenine methylase)^19,20^. We aimed to generate more robust datasets, both to benchmark existing callers, and to train and develop better methods for 6mA identification. To this end, we harnessed the methylation diversity in bacterial systems, which occurs in a highly motif-driven manner^22^. To create datasets that are methylated and unmethylated at specific sequence contexts, we sequenced both the native (NAT) and whole genome amplified (WGA) versions of *E.coli* K12 MG1655, and *Helicobacter pylori* 26695 strains on the Oxford Nanopore R9.4.1 flowcells (Fig 1A).

**Figure 1:**
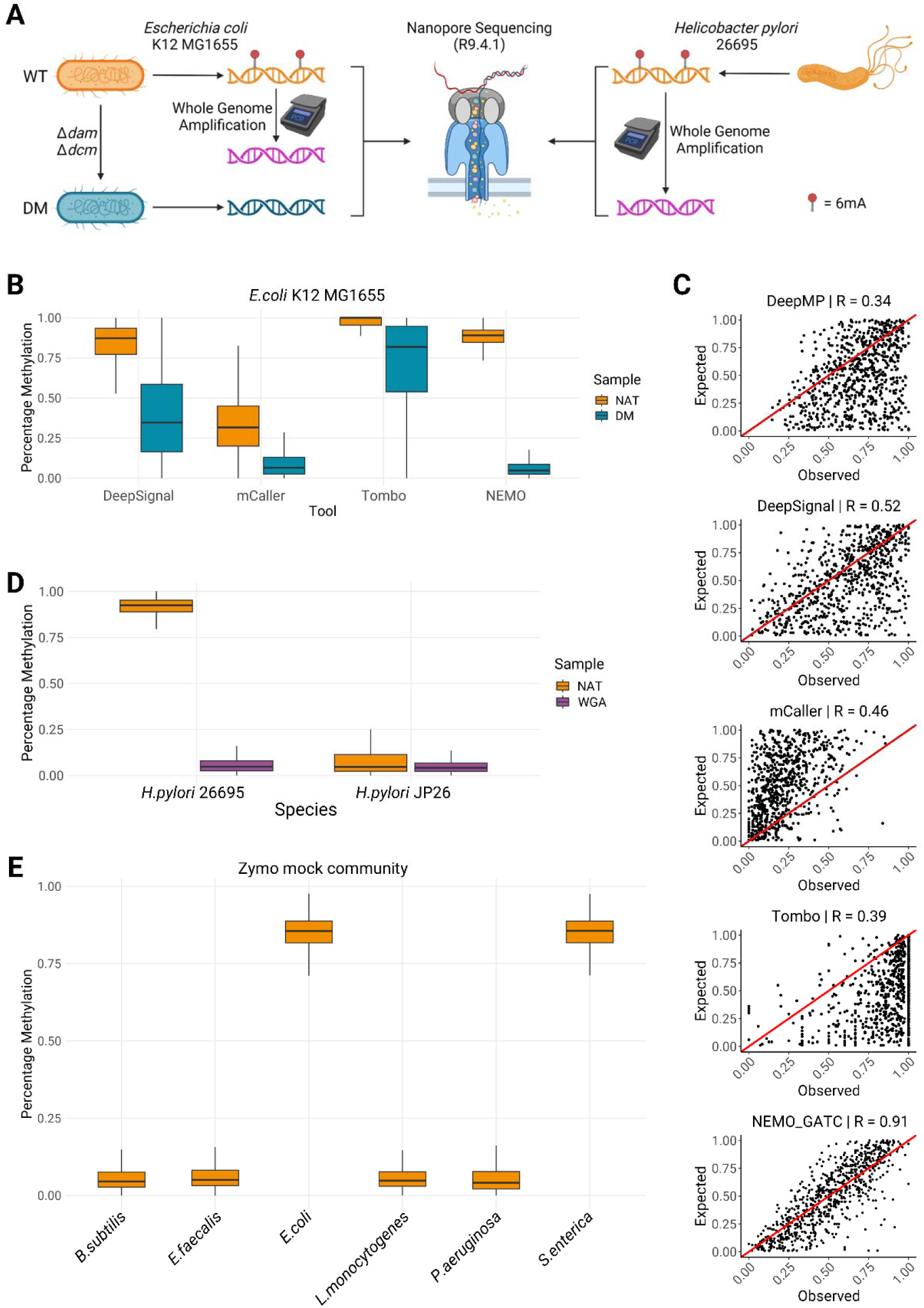
Performance of NEMO_R9_GATC. **A)** Schematic indicating the bacterial species and strains sequenced on Oxford Nanopore R9.4.1 chemistry to be used for model training and validation. NAT - Native, DM - double mutant, WGA - whole genome amplified. The schematic was created using BioRender (https://biorender.com). **B)** Performance comparison of NEMO_R9_GATC with other existing models on *E.coli* data. All GATC locations of the genome covered at least by 10 reads were profiled. **C)** Scatterplots comparing the performance of NEMO_R9_GATC with other tools on mixed data for 880 GATC sites in the *E.coli* genome. X-axis indicates the methylation percentage called by a tool whereas the Y-axis indicates the expected methylation. Red lines indicate the diagonal of complete concordance. Pearson correlation value is indicated in the plot title for each tool. **D)** Performance of NEMO_R9_GATC on native (NAT) and whole genome amplified (WGA) data of two different *Helicobacter pylori* strains, 26695 - where GATC methylation is present, and JP26 - where GATC methylation is absent. **E)** Performance of NEMO_R9_GATC on various bacterial species part of the Zymo mock community data taken from Sereika et al^24^. Of the 6 species represented here, GATC methylation is expected to be present in *Escherichia coli* and *Salmonella enterica*.

To control for possible amplification biases, we also created a “double mutant” (referred to as DM hereon) of *E.coli* K12 MG1655 substr. in which both the major methylases, dam and dcm, are deleted. We confirmed the insertion cassette and deletion in *dam* and *dcm* loci respectively in the DM strain using whole genome sequencing (Fig S1). Further, fragment analysis using TapeStation showed a characteristic smear for NAT genomic DNA (gDNA) digested with DpnI, which specifically cuts methylated GATC sites. However, DM gDNA was resistant to cleavage by DpnI, and showed an intact band comparable to undigested gDNA, indicating absence of methylation (Fig S2).

DpnI digested gDNA of both the NAT and DM strains were sequenced on the Oxford Nanopore platform. As expected, close to 100% of the reads spanning GATC sites in WT strain terminate at the cut site (GA/TC) whereas in DM, the read termination profile is similar to that of undigested gDNA (Fig S3). We derived the methylation percentage at GATC sites that were well covered by calculating the ratio of reads terminating at the cut site to the total number of reads mapped to the location. Our results indicate that all profiled GATC sites are close to fully methylated in the NAT strain, whereas completely unmethylated in the DM strain. This strain was also sequenced on R9.4.1 chemistry. In addition to the above, we also used data from two previous studies - native and WGA datasets from Tourancheau et al^23^, and the Zymo mock bacterial community data from Sereika et al^24^, to test the efficacy of our models in accurately identifying 6-methyladenine. The datasets used, and the sequence contexts tested in this work are summarized in Table 1.

### 6mA identification in GATC context

We first set out to develop a model that can discriminate 6mA from canonical adenine in GATC context (see Methods). We compared the performance of our GATC model (NEMO_R9_GATC) to existing tools. There are currently four different 6mA callers available - Tombo, DeepSignal, mCaller, and DeepMP. Of the 4, DeepSignal and DeepMP only support 6mA calling in a GATC context. Tombo and DeepSignal identified the highest number of methylated sites but included many false positives (Fig 1B). mCaller on the other hand missed many methylated sites but had fewer false positives. NEMO_R9_GATC outperformed all the other tools in identifying methylated sites with the least false positives (Fig 1B).

To test the efficacy of our model across the range of expected methylation values rather than 100% methylated or unmethylated sites, we generated a synthetic dataset by mixing known proportions of methylated and unmethylated reads for 880 GATC sites spanning the entire *E.coli* genome (See Methods). NEMO_R9_GATC showed a concordance of 0.91 on this dataset compared to the next best concordance of 0.52 by DeepSignal (Fig 1C). We did not see a difference in identification of unmethylated Adenine from both the DM data, as well as the whole genome amplified data of *E.coli* (Fig S4).

We further validated our model on datasets from multiple bacterial species to rule out any species- or lab-specific bias. To this end, we first analyzed data generated in-house from *H.pylori* 26695, where GATC methylation is expected, and data of another strain of *H.pylori,* JP26, from Tourancheau et al^23^, where GATC methylation was not found. NEMO_R9_GATC accurately discriminated 6mA and unmethylated Adenine in these datasets (Fig 1D). In addition, we tested it on previously published data of other bacterial species^22^. In all cases, our model reliably detected presence or absence of 6mA in GATC context in the respective genomes (Fig 1E). Taken together, these results indicate that the GATC model of NEMO for the R9.4.1 chemistry is highly accurate in discriminating 6mA from a canonical adenine.

### An all context 6mA model for R9 chemistry

Using native and WGA datasets of *H. pylori* 26695, we next aimed to train a model that could identify 6mA irrespective of the sequence context. The sensing regions of the R9.4.1 version of flow cells can harbor 5-6nt of DNA. Hence, depending on the sequence context, the signal to discriminate between canonical and modified bases can span up to 11-13nt, with the modified base at the center. With a sampling rate of 4kHz and translocation speed of 400 bases per second, this corresponds to approximately 130 signal points. We therefore tested multiple signal chunk sizes (30, 50, 75, 100, 120 and 150) around the target base to identify the optimal chunk size. We observed that the true positive calls improved with longer signal sizes, plateauing around 75 signal points, however, the true negative calling deteriorated at longer sizes, starting at around 75 signal points (Fig S5). In the contexts tested, the model trained with a chunk size of 50 data points (sig50) on either side of the target base performed most optimally. We further refer to the sig50 model as NEMO_R9_6mA.

We tested the efficacy of the NEMO_R9_6mA on *E.coli* K12 MG1655 and *H.pylori* 26695 data in diverse sequence contexts - motifs known to be generally methylated in the genome, and few other motifs where no methylation is expected. As seen in Fig 2A, our model performed well in discriminating 6mA from canonical adenine in the tested contexts on *H.pylori* data. When tested on *E.coli* data in ‘NANN’ context (all possible tetramers with adenine in the second position), NEMO_R9_6mA identified 6mA only in GATC context and in the native dataset, where methylation is expected (Fig S6). We then assayed the performance of NEMO_R9_6mA on different species using the Zymo mock community dataset from Sereika et al^24^. While the performance varied based on the sequence context, in all cases there was a clear difference in percentage methylated identified by NEMO in genomes where the motif is expected to be methylated, except for the motif CAGaG in *Salmonella enterica* (Fig 2B). When tested on synthetic data with methylated and unmethylated reads mixed in known proportions, the concordance for most tested contexts was >0.7 (Fig 2C, Fig S7). Lower concordance of ∼0.5 in contexts such as GaGG could be attributed to poor signal differences between the methylated and unmethylated samples, similar to what was observed for contexts where no methylation is expected (Fig S8).

**Figure 2:**
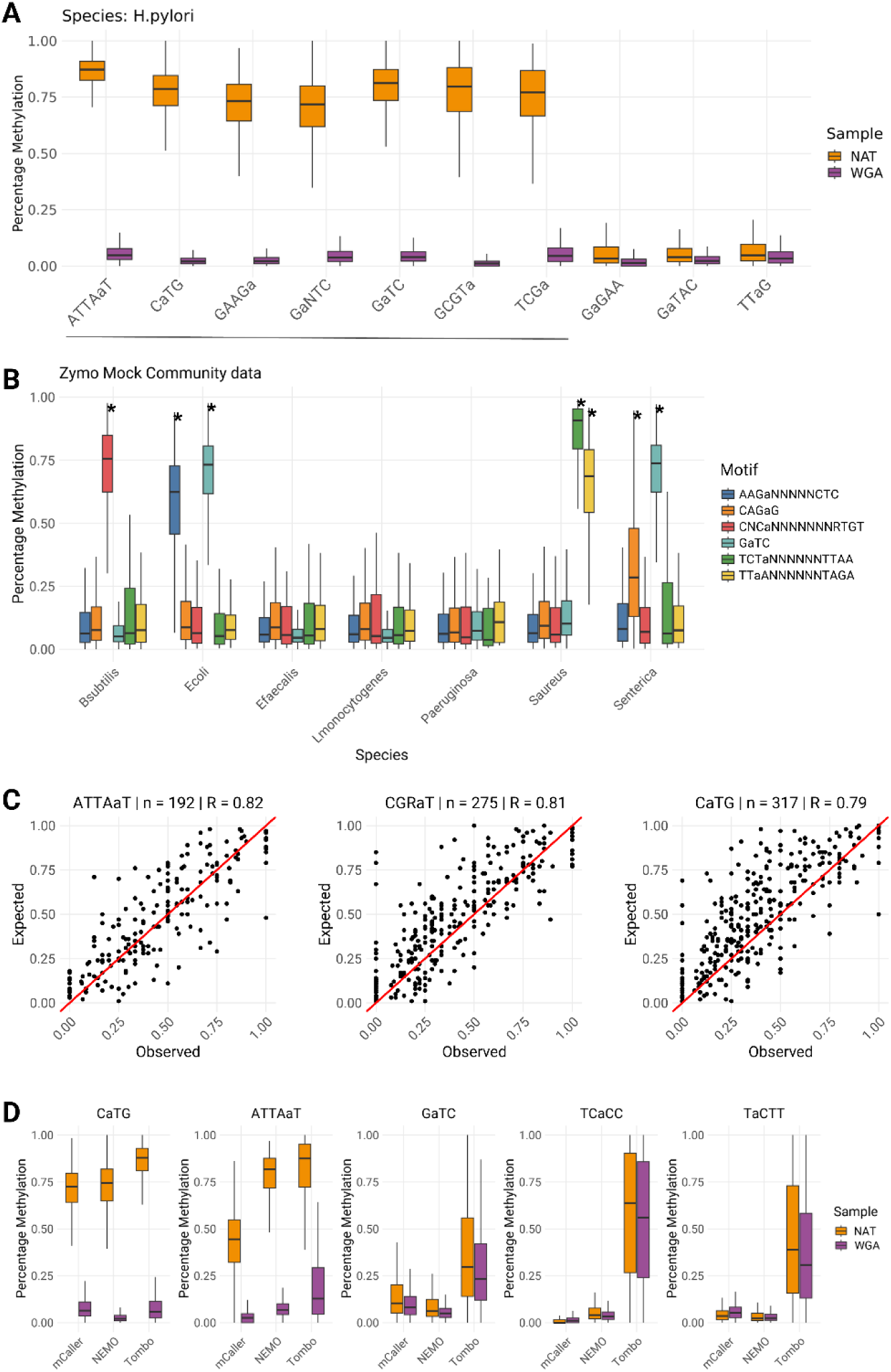
Performance of NEMO_R9_6mA. **A)** Evaluation of NEMO_R9_6mA on diverse sequence context using native (NAT) and whole genome amplified (WGA) data of Helicobacter pylori 26695. The first 7 (underlined) motifs are contexts where methylation is expected to be present, whereas the next 3 are not known to be methylated. **B)** Performance of NEMO_R9_6mA in identification of 6mA in various sequence contexts from diverse bacterial species from the Zymo mock community data. Motifs known to be generally methylated in a species are indicated with an asterisk. **C)** Scatterplots depicting the correlation between methylation values called by NEMO_R9_6mA and the expected ground truth. The motif, total number of genome locations profiled, and the Pearson correlation is indicated in the plot title. **D)** Performance comparison of NEMO_R9_6mA, Tombo, and mCaller in various sequence contexts, evaluated on the native (NAT) and whole genome amplified (WGA) data of *H.pylori* JP26 taken from Tourancheau et al^23^. In all plots, the adenine which is profiled for methylation is indicated in lower case in the motif.

We next wanted to compare NEMO_R9_6mA with the other two tools which support all context 6mA calling - Tombo and mCaller. Both Tombo and mCaller require the raw files to be preprocessed with the ‘resquiggle’ command of Tombo. However, Tombo does not support resquiggling on the newer POD5 files, even when these are converted to FAST5 format. Due to this incompatibility, we could not perform the comparison analysis on the in-house generated *H.pylori* 26695 data. Instead we used *H.pylori* JP26 data from Tourancheau et al^23^, which was in the older FAST5 format compatible with Tombo resquiggle. As was seen in the GATC context, in general Tombo overpredicted methylation, even in contexts where no methylation was expected (Fig 2D, Fig S9). mCaller performed well in discriminating 6mA in several contexts, particularly in identifying unmethylated adenines. However, it did not identify methylated adenine as effectively as Tombo and NEMO in a few sequence contexts such as ATTAaT and CATGa (Fig 2D, Fig S9).

We further profiled the performance of these tools in two motifs - aGGcC and aCcGG, where no adenine methylation is expected, but there is cytosine methylation (5mC and 4mC respectively). In these cases, mCaller identified many false positives for 6mA in the context aGGcC but not aCcGG (Fig S9), suggesting that its model is sensitive to neighboring 5-methylcytosine but not 4-methylcytosine. In a few cases such as GaATTC, NEMO did not perform as well compared to Tombo in calling methylated adenine. Overall, our results show that NEMO_R9_6mA is able to reliably detect 6mA in a variety of sequence contexts from nanopore data generated on the R9.4.1 chemistry.

## Discussion

The advent of long-read sequencing technology has opened new avenues for studying sequence and epigenetic variation simultaneously. The ability to analyze epigenomes concurrently with sequencing is contingent upon availability of robust models capable of accurately discriminating modified bases. The support to identify 6mA from nanopore data has been limited, particularly for data from the R9 versions of the flowcells, which have been one of the most widely used FC versions. For example, the 6mA models offered by Guppy and Dorado do not support methylation calling on data generated on flowcells prior to version R10.4.1. Previous tools that support 6mA calling have significant limitations in speed, accuracy, convenience, and support for new file formats. Our tool, NEMO, addresses these limitations.

We report two different models of NEMO - one specific for GATC context, and one which is sequence context-agnostic. In addition to studying bacterial genomes where GATC is prevalent, accurate identification of 6mA in GATC context has applications in techniques such as DamID^25^. Furthermore, 2 out of the 4 tools we used for performance comparison support 6mA identification in GATC context alone. Compared to other tools, NEMO achieves significantly higher concordance scores.

In our quest to develop a sequence context-agnostic model for 6mA identification, we standardized the signal chunk used in training. We found that a chunk size of 50 signal points is optimal. While true positive rates remain high with larger chunks, the false positive rate increases dramatically, likely due to the training dataset (*H. pylori*) having abundant 6mA, possibly leading to confusion in signal interpretation due to neighboring modifications. The other possible explanation is that the true signal that can discriminate 6mA is diluted by the noise introduced in a large signal chunk. Further research is needed to understand the performance decline with longer signal chunks.

Previous tools have enabled 6mA identification from nanopore data, but often as standalone applications. During our comparative experiments, we encountered challenges in running these tools on newer file formats, such as POD5. The resquiggle command of Tombo, a mandatory requirement for other tools, did not work on POD5 files, even when they were converted to FAST5 files using pod5tools. However, this gave us an opportunity to test the NEMO models on data available in the public domain, to ensure that they also work well on data generated beyond our own lab. Additionally, while some tools supported GPU acceleration, others did not, and each required its specific setup. In contrast, NEMO models seamlessly integrate with the latest base callers by Oxford Nanopore, such as Bonito and Dorado. NEMO does not require any specific setup and works out-of-the-box, leveraging any available GPU acceleration to significantly improve processing times. This not only boosts efficiency but also offers greater convenience to users.

Our “all-context” 6mA model does show limited capabilities in few contexts. This is perhaps due to the fact that the signal difference in these contexts could not be captured efficiently. This may be a limitation of the flow cell itself, where the shorter barrel of the R9 nanopore is not capable of discriminating 6mA in all sequence contexts. Alternatively, a model trained using a shorter signal chunk may bring out the weak signal differences and hence perform better in these contexts. In future, a consensus taken by using two different models, one trained on short chunks and one on long chunks, may provide a more wholesome understanding of 6mA in a dataset.

## Methods

### Bacterial strains and culture

The “double mutant” (DM) strain was constructed in the background of *Escherichia coli* MG1655. dam::Cm allele (dam locus harboring transposon Tn9; M.R. lab collection) and dcm::Kan^26^ were introduced singly or sequentially into MG1655 by P1 phage mediated transduction. Deletions were confirmed by PCR. Strains were grown at 37°C in lysogeny broth (1% tryptone, 0.5% yeast extract, 1% NaCl) supplemented with chloramphenicol (Cm; 10μg/ml) or kanamycin (Kan; 25μg/ml) as required.

### Genomic DNA extraction

An overnight bacterial culture was treated with Lysozyme and then lysed using SDS and Proteinase K. The sample was treated with CTAB to further purify the gDNA. This was followed by a Phenol:Chloroform:Isoamyl alcohol and Chloroform:Isoamyl alcohol wash. DNA was precipitated using Isopropanol and spooled onto a glass rod. The gDNA was washed in ethanol, air dried and finally dissolved in EB (Qiagen, Germany). The gDNA was stored at 4°C for 2 days before quantification using a NanoDrop 2000 Spectrophotometer (ThermoScientific, USA) and Qubit dsDNA kit (ThermoScientific, USA).

### *in vitro* DNA Methylation

Genomic DNA from DM strain was used for in-vitro DNA methylation. The genomic DNA was incubated in a mix containing S-Adenosyl methionine (SAM) along with the DNA adenine Methyltransferase (dam, NEB, USA) and its corresponding buffer for 4 hours at 37°C. The sample was purified using Ampure XP (Beckmann Coulter, USA) beads and the methylated DNA was quantified using Qubit HS dsDNA kit (ThermoScientific, USA).

### Nanopore library preparation

Nanopore library was constructed using the ligation sequencing kit SQK-LSK109 (Oxford Nanopore Technologies, UK). The DNA was treated with NEBNext Ultra II End Repair/dA-Tailing Module (NEB, USA) and NEBNext FFPE DNA Repair Mix (NEB, USA), purified using Ampure XP (Beckmann Coulter, USA) and quantified using Qubit dsDNA (ThermoScientific, USA). To multiplex the samples, the Native Barcoding Expansion kit (Oxford Nanopore Technologies, UK) was used. The barcodes were ligated to samples using the NEBNext Ultra II Ligation Module (NEB, USA) after which the samples were purified and quantified. Equal amounts of barcoded samples were pooled and ligated to the adaptor protein. The final library was loaded onto an R9.4.1 flow cell (Oxford Nanopore Technologies, UK) as per the manufacturer’s instructions.

### Nanopore Sequencing Data Processing

We used Guppy (V6.3.7) for basecalling all the nanopore reads (fast5 files) using super accuracy models (sup) with options ‘--q-value 7 --fast5_out’. Quality check for the basecalled reads (fastq files) was performed using NanoPlot^27^. The basecalled reads were mapped to the reference genome using minimap2^28^. We compared our model against 4 other existing 6mA calling tools - DeepSignal, mCaller, Tombo, DeepMP. DeepSignal v0.1.6 was run using the GATC model with options ‘--motifs GATC --mod_loc 1’. Tombo v1.5.1 was run with the alternative model using options ‘--alternate-bases 6mA & --alternate-bases dam’ independently to be able to predict 6mA methylation in individual samples. mCaller v1.0.3 was run using model ‘r94_model_NN_6_m6A.pkl’ with parameters ‘-m A -b A’. We used DeepMP for combined feature extraction followed by modification calling using pre-trained model ‘pUC19_joint_202106’ for 6mA detection in GATC context. The snakemake files for these tools were adapted from the METEORE tool (https://github.com/comprna/METEORE).

### Restriction digestion based analysis

To validate the presence or absence of methylation in GATC context, NAT, DM and DM+DAM treated gDNA was digested with DpnI (NEB, USA), which selectively cleaves methylated GATC sites. Digested gDNA and undigested controls were sequenced on the Oxford Nanopore platform. The sequenced data was aligned to the *E.coli* reference genome using minimap2, and the bam files were analyzed using a custom python script which leverages the pysam library (https://github.com/pysam-developers/pysam) to determine the frequency of reads that terminate or start at GATC sites; The indexed bam files are parsed using pysam and the read positions are then iterated over using the ‘pileup’ method. Each of the pileups-iterable objects can then be tested for read termination or initiation using the ‘is_tail’ and ‘is_head’ methods respectively. The sum of these individual values indicate the frequency of reads that terminate or start at a given genomic location. The ratio of the number of read terminations/start-sites to the sequencing depth at a given location is used to calculate the percentage methylation at a given GATC site. Data was visualized using IGV and ggplot2.

### Model training in GATC context

Custom models were built using remora v0.2.4 (https://github.com/nanoporetech/remora). First, we used the ‘dataset prepare’ function to get motif specific chunks of signal data from the training datasets (POD5 files). We used DM as the canonical dataset and NAT as the modified dataset for model training in GATC context with parameter ‘--motif GATC 1’. The motif specific chunks of signal from DM & NAT were then merged and given as input for model train function of remora. The output was the directory containing the best model in PyTorch format.

### All context 6mA model training

An all context model for 6mA detection was also trained using remora v0.2.4 (https://github.com/nanoporetech/remora). Native genomic DNA and WGA-amplified DNA samples for *Helicobacter pylori* 26695 were sequenced using R9.4.1 flowcells (Oxford Nanopore Technologies, UK). We used the whole genome amplified sample (WGA) as canonical dataset and native sample (NAT) as the modified dataset for training an all context model with parameters ‘--motif A 0’, ‘--refine-rough-rescale’.

### Optimization of the neural network architecture

The basic layout of the neural network is a convolutional long-short-term-memory (LSTM) model. To ensure the NEMO models are accurate and not overfit, we optimized various parameters of the neural network, including the chunk sizes that were used for training. We trained the models from 10 to 100 epochs, and pool layer sizes of 64-256. The sensing regions of the R9.4.1 version of flow cells can harbor 5-6nt of DNA at a time. Therefore, the location of the signal to discriminate between canonical and methylated adenines can be different based on the motif. For the chunk context parameter, we used multiple signal chunk sizes (30, 50, 75, 100, 120 and 150) around the target base to generate different models to look at the effect of signal chunk size on the accuracy of the model. Overall, the optimization effort took over 4000 GPU hours. Finally, we used a layer size of 64 and a chunk size of 50 for NEMO_R9_GATC, and layer size of 256 and a chunk size of 50 for NEMO_R9_6mA. The best performing model in each case was exported as a PyTorch and dorado models using remora, which are compatible with bonito and dorado basecallers respectively.

### Model inference

To assess the efficacy of the models, we used in-house datasets generated for *E.coli* K12 MG1655 and *Helicobacter pylori* 26695 strains on R9.4.1 flowcells (Oxford Nanopore Technologies, UK). In addition, we also used data from two previous studies, Tourancheau et al^23^, and Sereika et al^24^. We used dorado v0.3.3 (https://github.com/nanoporetech/dorado) with a super accuracy model and NEMO model for performing modified basecalling on above mentioned datasets. Dorado outputs reference aligned reads with the methylation information in modbam format. The modbam file was then sorted and indexed using samtools. Further, to aggregate modified base counts stored in modbam file, modbam2bed tool (https://github.com/epi2me-labs/modbam2bed) was used with the parameter “-m 6mA”. The bed file containing methylation calls was processed for data wrangling and visualization using the R packages dplyr and ggplot2.

### Simulated data with known methylation levels

To benchmark the performance of NEMO at different methylation levels we created a dataset with known methylation levels at the sequence context of interest. The pod5 files for WGA and NAT runs for a specific species were first basecalled to FASTQ files independently, using dorado. These reads are then size selected using NanoFilt (https://github.com/wdecoster/nanofilt) for a minimal size of 500bp and a maximum length of 4 kb. Once a motif of interest is determined, all corresponding motif locations across the genome are identified using an inhouse script. These motifs are then filtered such that they are at least 5 kb apart (1kb + max size of reads), in order to eliminate reads spanning multiple motifs and thus interfering with the methylation percentage at the time of mixing. Once a list of motif sites are determined, a randomizer is used to generate an arbitrary percentage of methylation for each site, with a constant seed value for consistency across multiple mixing experiments performed using different sample combinations and motifs.

The FASTQ reads are then aligned to the respective reference genomes using minimap2, thus generating two files one for the reads containing no methylated sites for the motif of interest (canonBam) and another with the reads that contain 6mA nucleotides in the motif of interest (modBam). After alignment the bam files are converted to a bed format for both canonical and modified data. These coordinate bed files are filtered for reads that span the filtered motif list that was generated earlier. Reads from each bed file are combined in appropriate combinations based on the methylation percentages generated and the final output is a TSV file containing readID and a second column describing the origin of the read (canon/mod). Using the subset utility in the POD5 python package (https://pypi.org/project/pod5/), along with the original POD5, we created subsetted POD5 files with known methylation levels from original POD5 files. These were then used to infer the accuracy of various models. For NEMO model accuracy, we used bonito basecaller (https://github.com/nanoporetech/bonito) for methylation calling.

### Significance testing for sequence contexts

The Remora API was utilized to extract raw signal data from POD5 files from 100 randomly selected genomic locations for each sequence context. This was done for both NAT and WGA datasets of *Helicobacter pylori* 26695. Signal was trimmed to 20 base positions, with 10 bases before and after Adenine. We calculated metric values such as dwell, trimmean and trimsd for each of the 100 locations, using a maximum of 1000 reads per location. We then calculated p values for each base position by performing a t-test on trimmean metric of signals from NAT and WGA datasets. These p values were plotted using ggplot2 R package.

## Ethics approval and consent to participate

Not applicable

## Consent for publications

Not applicable

## Availability of data and methods

The datasets generated as part of this work are deposited to the NCBI Sequence Read Archive under the PRJNA1076669. NEMO models are available in our github repository at https://github.com/SowpatiLab/NEMO.

## Author contributions

Conception and design of the study: DTS, KBT, MR; Data acquisition: LZ, SG, SA; Data analysis and interpretation: OK, RJM, NKS; Manuscript preparation: DTS, OK, RJM, LZ; Manuscript editing: MR, KBT; Funding acquisition: KBT, DTS. All authors have read and approved the manuscript.

## Acknowledgements

We would like to thank Dr. Santosh Kumar and Apoorva Etikala for the genomic DNA of *Helicobacter pylori* 26695. The schematic in Figure 1 was created using BioRender (https://biorender.com).

## Figure legends

**Fig S1:** IGV screenshots depicting the insertion disrupting the dam locus (top) and the deletion at dcm locus (bottom) of *Escherichia coli*. Red - Native (Wildtype) strain, Blue - Double Mutant.

**Fig S2:** Fragment analysis using TapeStation showing sensitivity or resistance of genomic DNA to DpnI cleavage. DpnI specifically cuts methylated GATC sites. Undigested samples show intact DNA with molecular weight >15-20kb (1 replicate each). gDNA from Native *E.coli* is sensitive to DpnI cleavage as indicated by the smear, whereas gDNA from Double Mutant is comparable to Undigested samples. Introduction of GATC methylation using in vitro treatment with DAM methylase renders DM gDNA susceptible to DpnI.

**Fig S3:** IGV screenshot showing the read termination profiles of various *E.coli* strains. gDNA that is cleaved by DpnI shows that all reads terminate at GATC, whereas other samples show random termination profile, comparable to undigested DNA.

**Fig S4**: Performance of NEMO_R9_GATC on various data of *E.coli*. Native *E.coli* data (NAT) shows close to 100% methylation, whereas both double mutant (DM) and whole genome amplified (WGA) data show close to 0% methylation, indicating no PCR bias in model performance.

**Fig S5**: Effect of signal chunk size on the accuracy of NEMO models in 6mA identification. For motifs where methylation is expected (Top row), accuracy begins to plateau at signal size of 75. Similarly, the accuracy of negative prediction in motifs where no methylation is expected (Bottom row) begins to drop at signal size of 75.

**Fig S6**: Performance of NEMO_R9_6mA on all tetramers with the profiled adenine at the second position, on native (orange) and whole genome amplified (purple) data of *Escherichia coli*. Methylation is only expected in the sequence context GaTC. The all context model of 6mA does well in discriminating 6mA from canonical adenine in all tested tetramer contexts.

**Fig S7**: Scatterplots depicting the correlation between methylation values called by NEMO_R9_6mA and the expected ground truth. The motif, total number of genome locations profiled, and the Pearson correlation is indicated in the plot title. The adenine which is profiled for methylation is indicated in lower case.

**Fig S8:** Signal differences between canonical and modified adenines in various sequence contexts. For each sequence context, the line plot on the right indicates statistical significance of signal differences between canonical and modified bases from 100 loci, for -10nt to +10nt around the target adenine. The performance of NEMO_R9_6mA for that sequence context is shown on the left. The six motifs chosen are based on the expected methylation - methylation expected: ATTAaT, CaTG, GCGTa, GaGG, methylation not expected: GaGAA, TTaAG. The profiled adenine is indicated in lowercase in the motif.

**Fig S9:** Performance comparison of NEMO_R9_6mA, Tombo, and mCaller in various sequence contexts, evaluated on the native (NAT) and whole genome amplified (WGA) data of *H.pylori* JP26 taken from Tourancheau et al^23^. In all plots, the adenine which is profiled for methylation is indicated in lower case in the motif. The motifs GGCC and CCGG are expected to show Cytosine methylation (5mC and 4mC respectively).

