## Supplementary Figures for "NEMO: Improved and accurate models for identification of 6mA using Nanopore sequencing"

### Fig S1

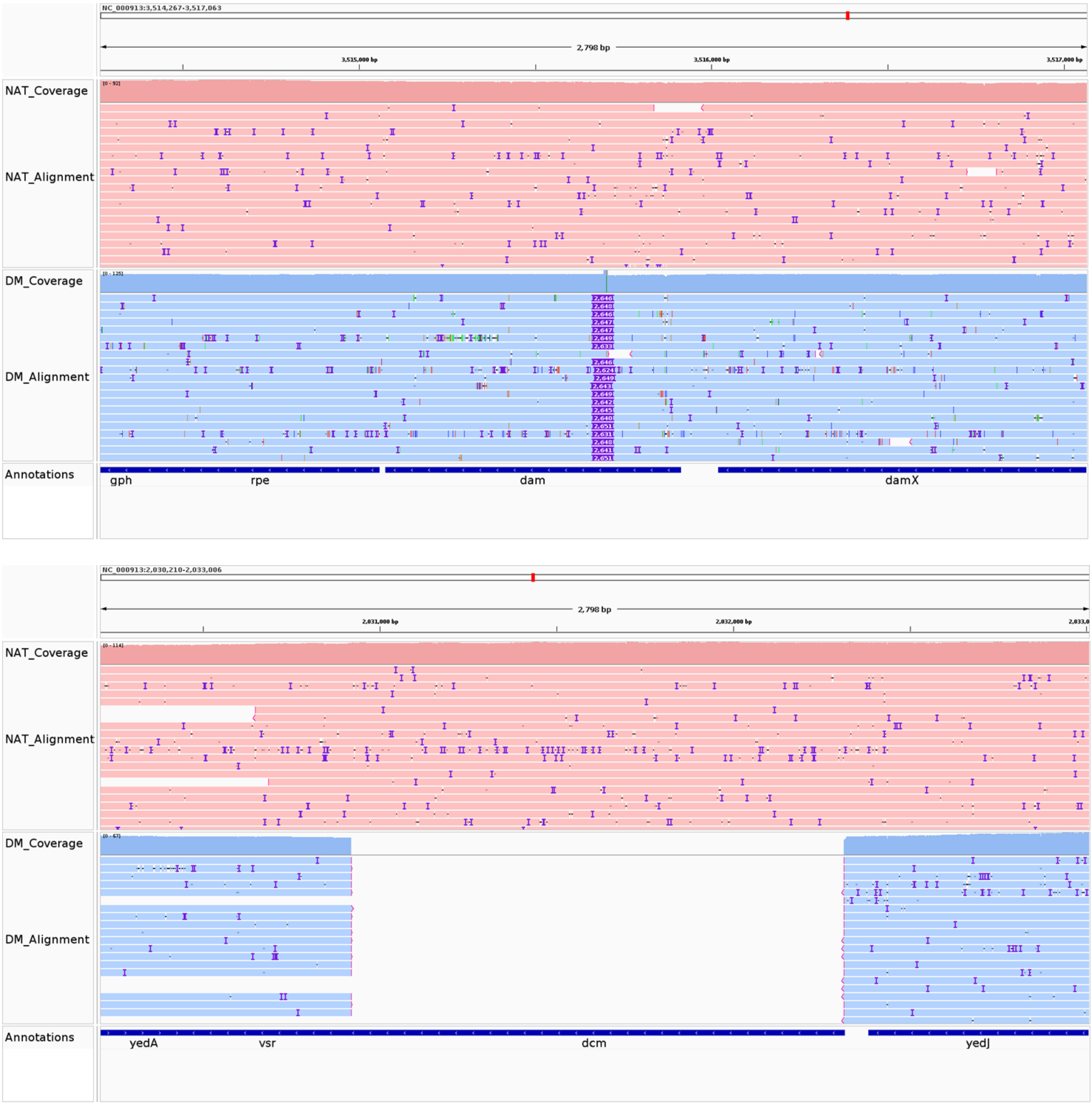


**Fig S1:** IGV screenshots depicting the insertion disrupting the dam locus (top) and the deletion at dcm locus (bottom) of *Escherichia coli*. Red - Native (Wildtype) strain, Blue - Double Mutant.

### Fig S2

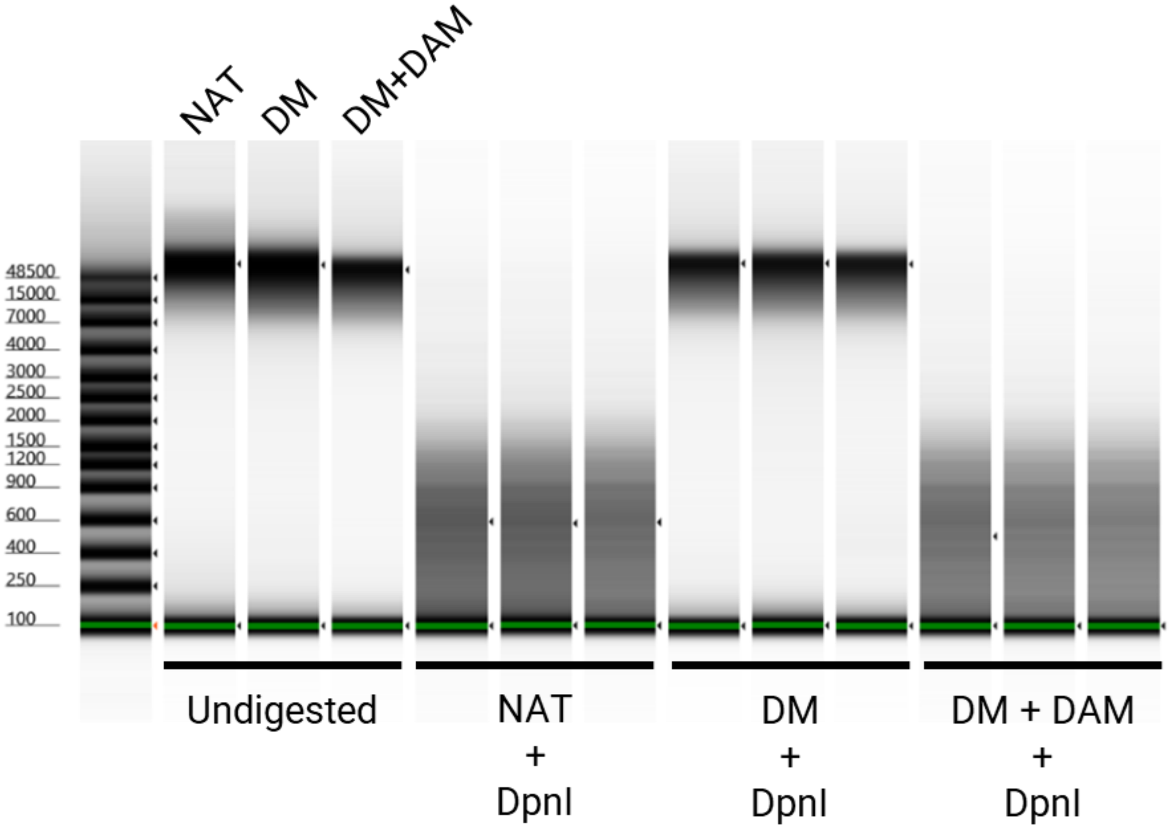


**Fig S2:** Fragment analysis using TapeStation showing sensitivity or resistance of genomic DNA to DpnI cleavage. DpnI specifically cuts methylated GATC sites. Undigested samples show intact DNA with molecular weight >15-20kb (1 replicate each). gDNA from Native *E.coli* is sensitive to DpnI cleavage as indicated by the smear, whereas gDNA from Double Mutant is comparable to Undigested samples. Introduction of GATC methylation using in vitro treatment with DAM methylase renders DM gDNA susceptible to DpnI.

### Fig S3

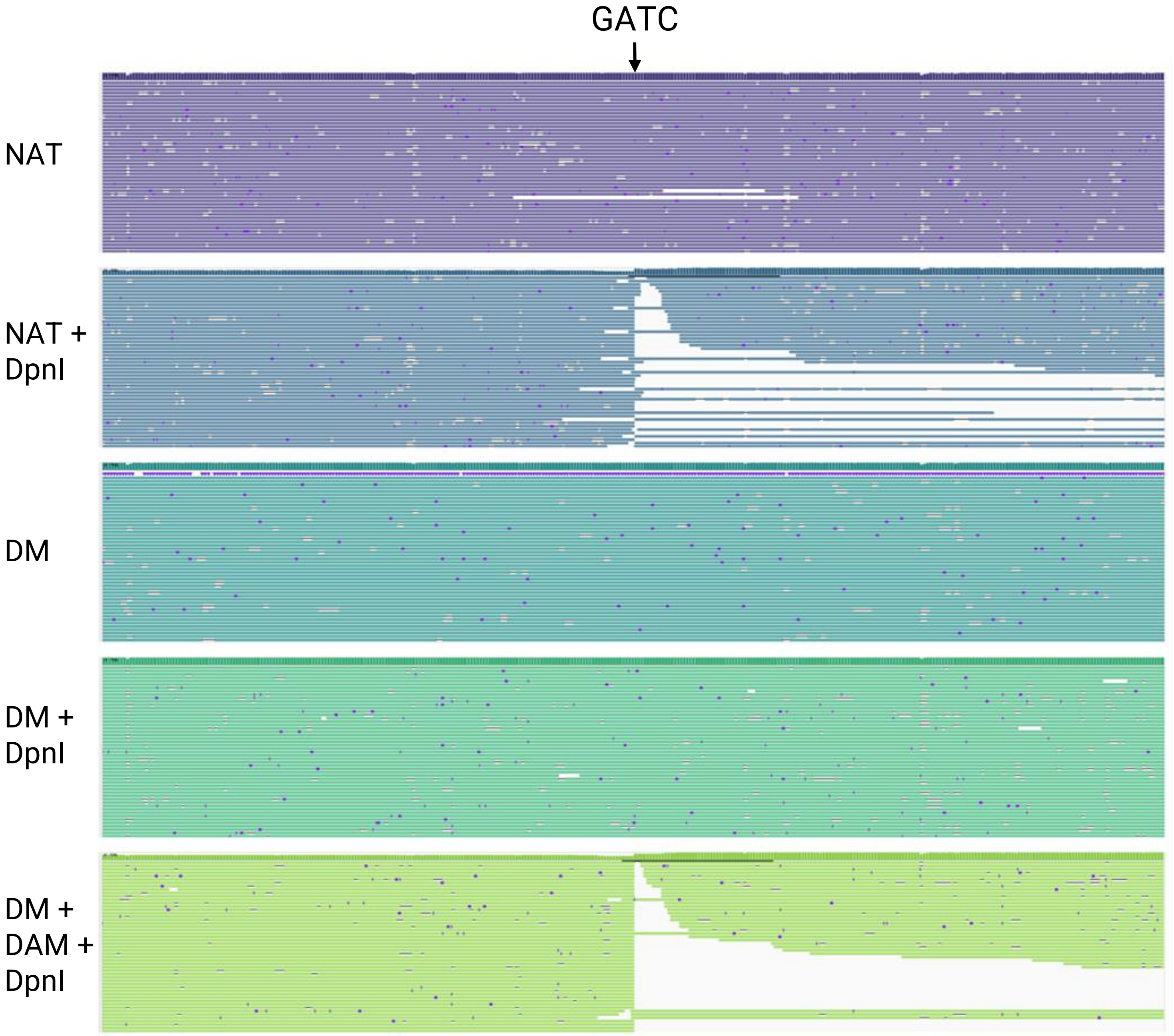


**Fig S3:** IGV screenshot showing the read termination profiles of various *E.coli* strains. gDNA that is cleaved by DpnI shows that all reads terminate at GATC, whereas other samples show random termination profile, comparable to undigested DNA.

### Fig S4

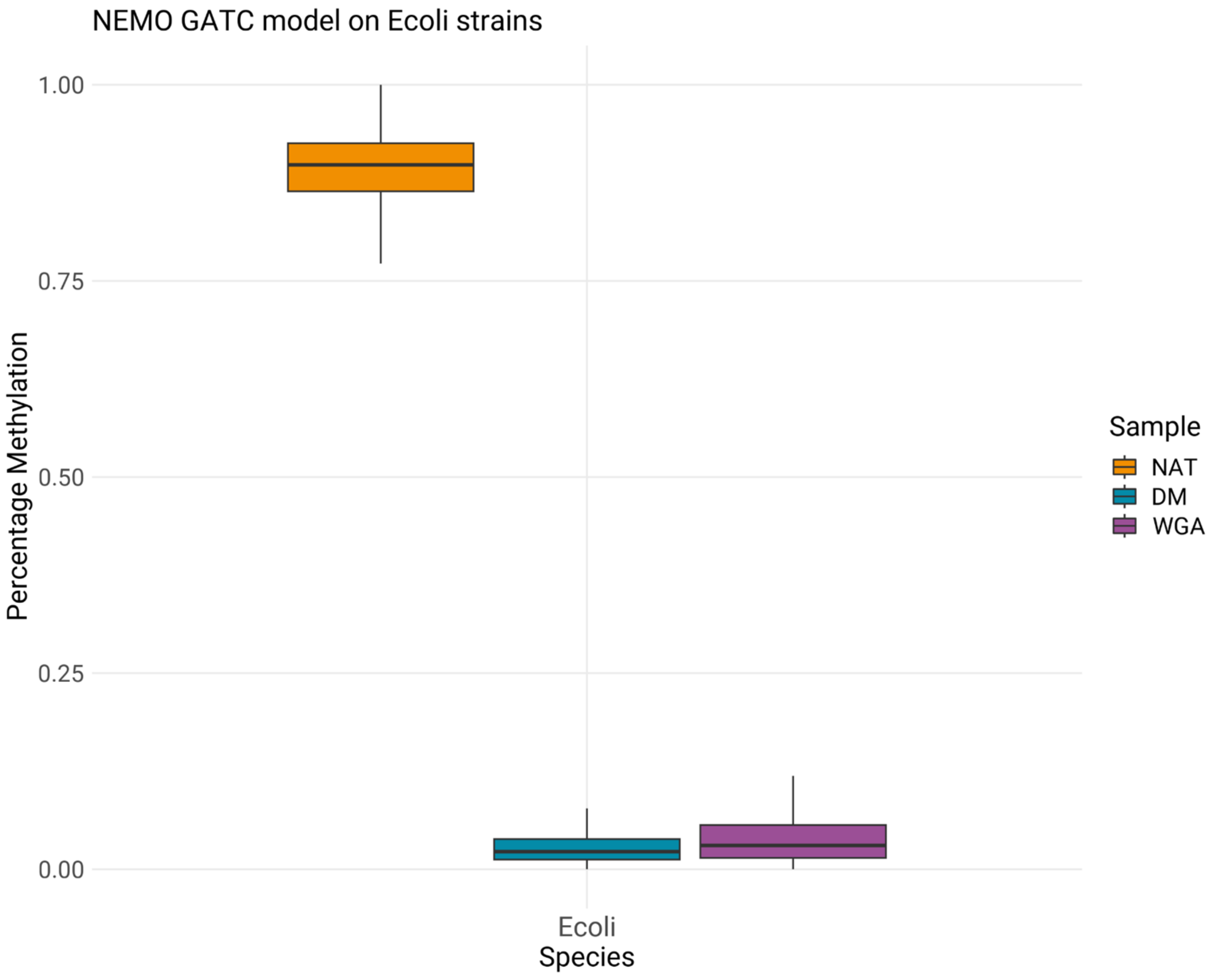


**Fig S4:** Performance of NEMO_R9_GATC on various data of *E.coli*. Native *E.coli* data (NAT) shows close to 100% methylation, whereas both double mutant (DM) and whole genome amplified (WGA) data show close to 0% methylation, indicating no PCR bias in model performance.

### Fig S5

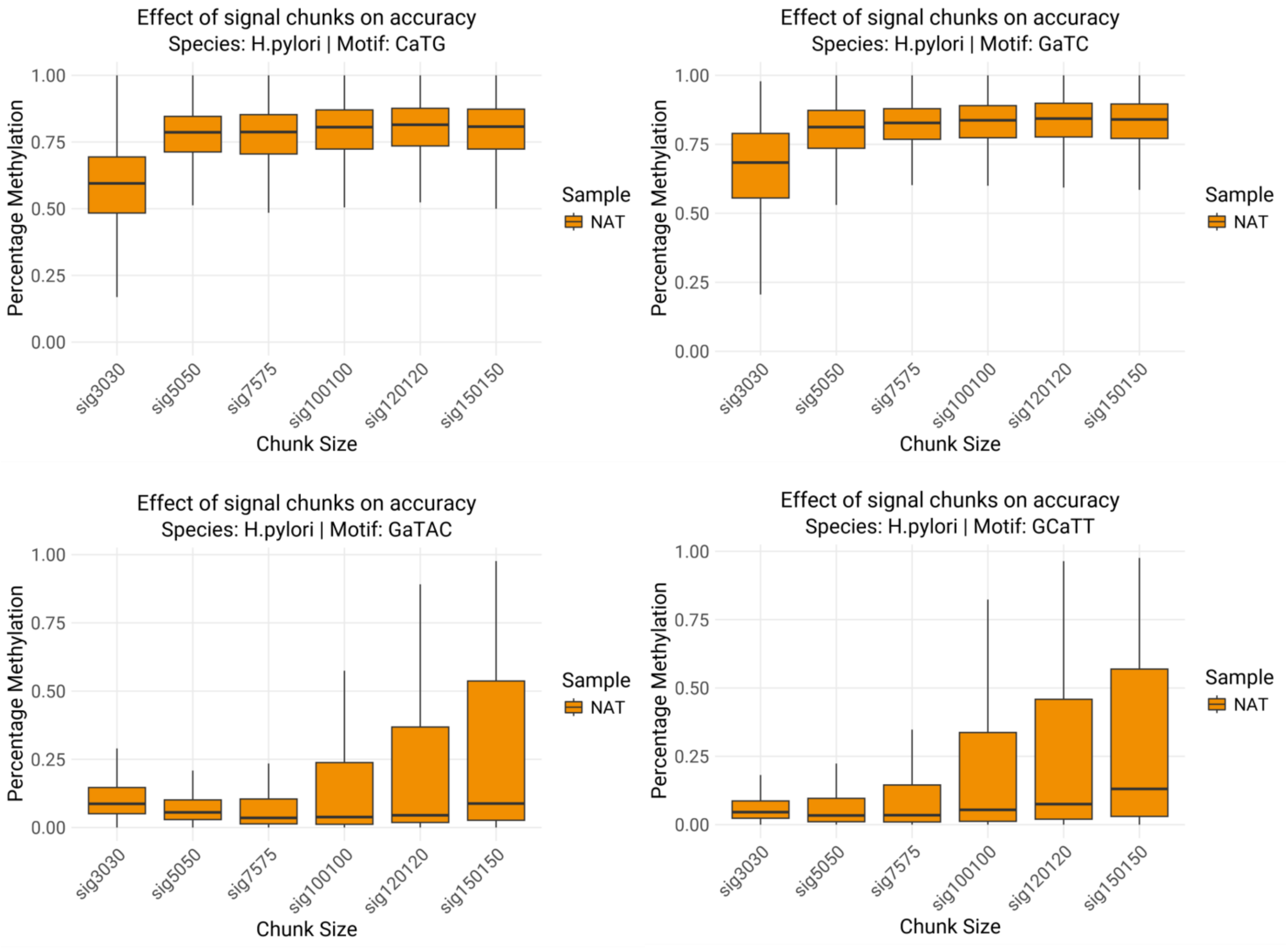


**Fig S5:** Effect of signal chunk size on the accuracy of NEMO models in 6mA identification. For motifs where methylation is expected (Top row), accuracy begins to plateau at signal size of 75. Similarly, the accuracy of negative prediction in motifs where no methylation is expected (Bottom row) begins to drop at signal size of 75.

### Fig S6

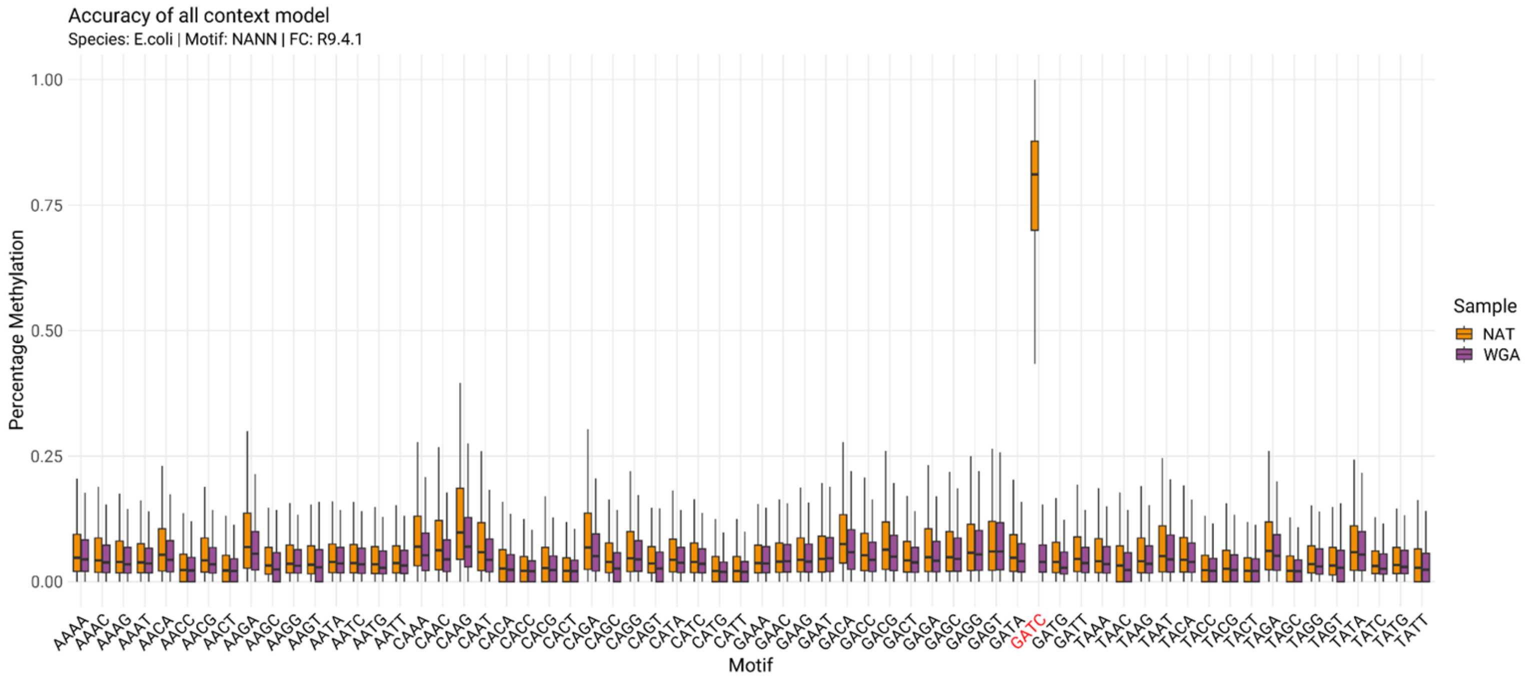


**Fig S6:** Performance of NEMO_R9_6mA on all tetramers with the profiled adenine at the second position, on native (orange) and whole genome amplified (purple) data of *Escherichia coli*. Methylation is only expected in the sequence context GaTC. The all context model of 6mA does well in discriminating 6mA from canonical adenine in all tested tetramer contexts.

### Fig S7

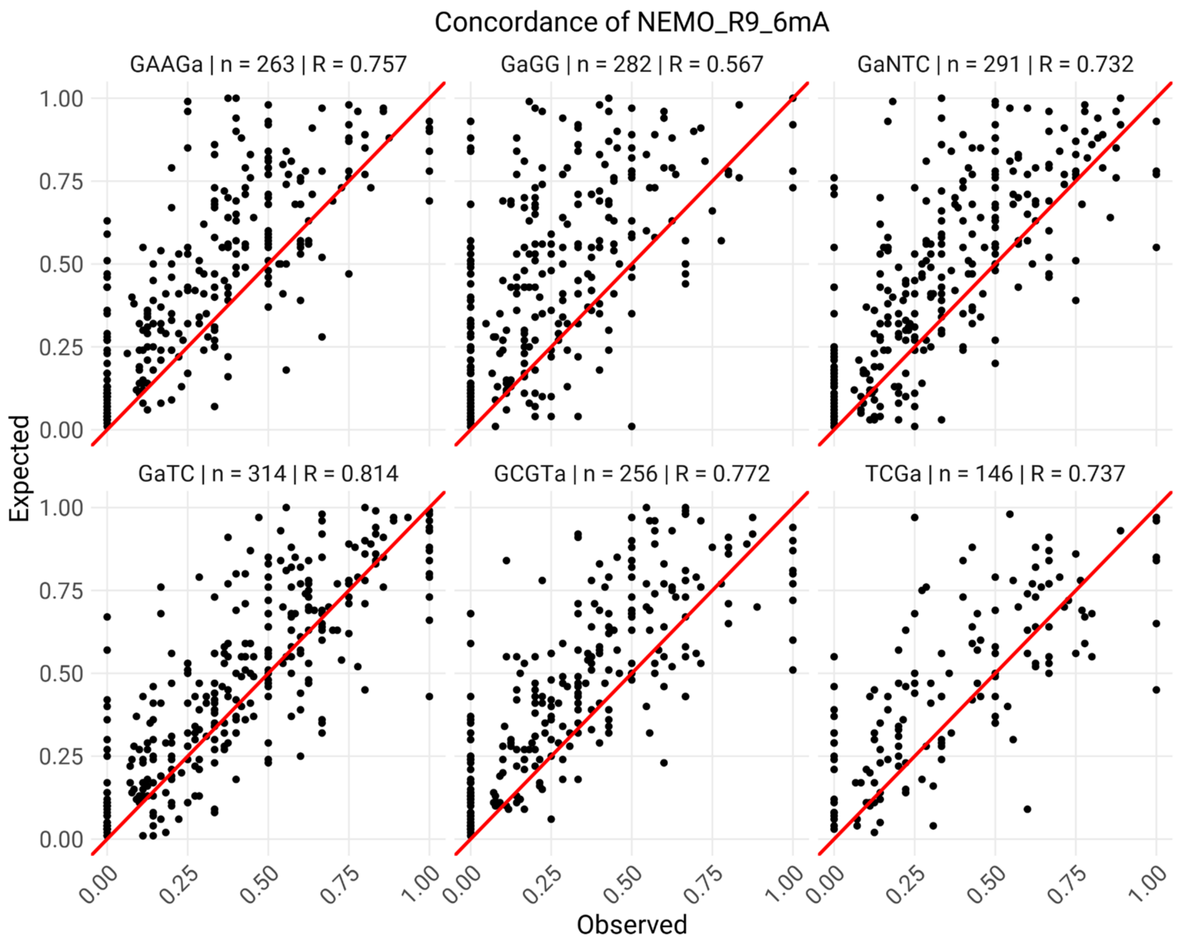


**Fig S7:** Scatterplots depicting the correlation between methylation values called by NEMO_R9_6mA and the expected ground truth. The motif, total number of genome locations profiled, and the Pearson correlation is indicated in the plot title. The adenine which is profiled for methylation is indicated in lower case.

### Fig S8

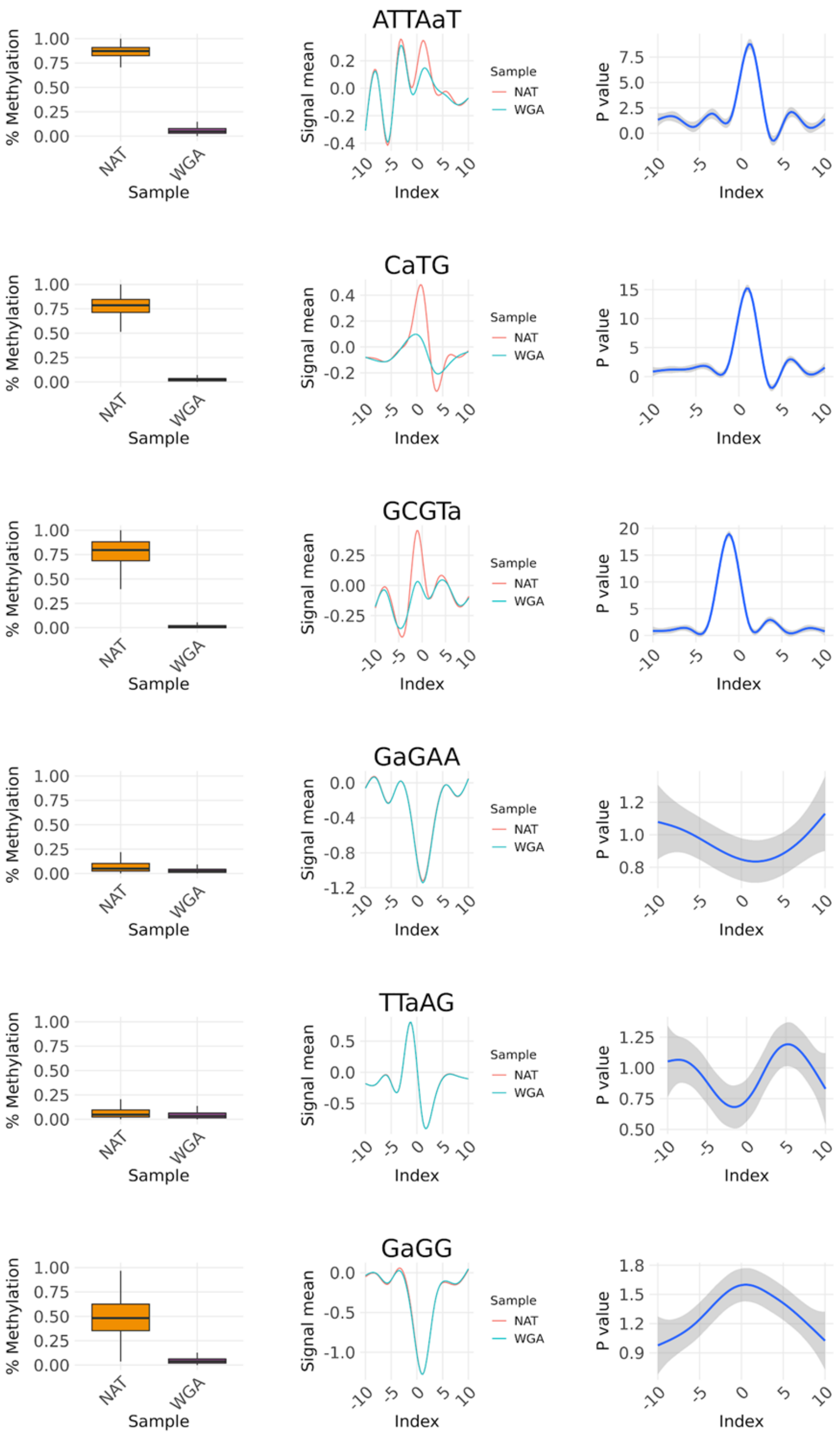


**Fig S8**: Signal differences between canonical and modified adenines in various sequence contexts. For each sequence context, the line plots in the middle indicate the signal means compared to baseline, on the right indicates statistical significance of signal differences between canonical and modified bases from 100 different loci, for -10nt to +10nt around the target adenine. The performance of NEMO_R9_6mA for that sequence context is shown on the left. The six motifs chosen are based on the expected methylation - methylation expected: ATTAaT, CaTG, GCGTa, GaGG, methylation not expected: GaGAA, TTaAG. The profiled adenine is indicated in lowercase in the motif.

### Fig S9

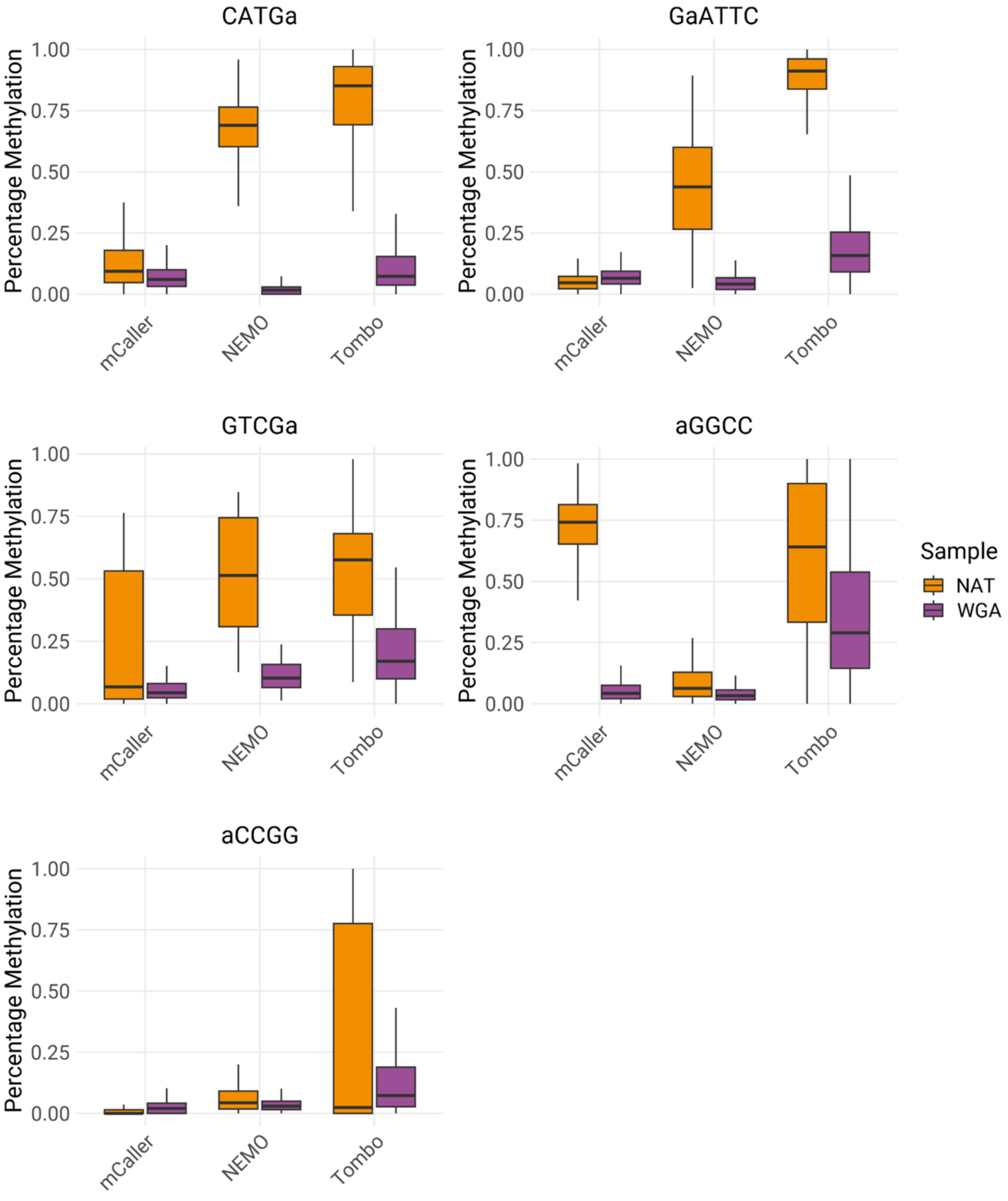


**Fig S9**: Performance comparison of NEMO_R9_6mA, Tombo, and mCaller in various sequence contexts, evaluated on the native (NAT) and whole genome amplified (WGA) data of *H.pylori* JP26 taken from Tourancheau et al[23]. In all plots, the adenine which is profiled for methylation is indicated in lower case in the motif. The motifs GGCC and CCGG are expected to show Cytosine methylation (5mC and 4mC respectively).
